# Disruptive selection via pollinators and seed predators on the height of flowers in a wind-dispersed alpine herb

**DOI:** 10.1101/2022.06.25.497599

**Authors:** Kai-Hsiu Chen, John R. Pannell

## Abstract

**Premise of the study:** Floral stalk height is known to affect pollination and seed dispersal in wind-dispersed grassland species, but it may also affect the attractiveness of flowers and fruits in animal-pollinated and animal-dispersed plants. Stalk height may thus be responsive to selection via interactions with both mutualist pollinators and seed dispersers, but also antagonist florivors and seed predators. In this study, we aimed to determine the effect of pollinators and seed predators on selection on floral stalk height in the insect-pollinated and wind-dispersed alpine andromonoecious herb *Pulsatilla alpina*, whose flowers also vary in their sex allocation and thus in the resources available to both mutualists and antagonists.

**Methods:** We measured the resource status of individuals in terms of their size and the height of the vegetation surrounding plants of *P. alpina* at eleven sites. In one population, we recorded the dynamics of floral stalk height over the course of an entire growing season and its association with floral morphology and floral sex allocation (pistil and stamen number), and we used leaf-removal manipulations to assess the effect of herbivory on floral stalk height. Finally, in four populations we studied phenotypic selection on floral stalk height in four female components of reproductive success before seed dispersal.

**Key results:** Stalk height was positively associated with female allocation of the respective flower, the resource status of the individual, and the height of the surrounding vegetation, and negatively affected by leaf removal. Our results point to disruptive selection on stalk height in terms of both selection differentials and selection gradients for fertilization, seed predation, and seed maturation rates, and to positive selection on stalk height in terms of a selection differential for mature seed number.

**Conclusions:** Stalk height in *P. alpina* is a costly trait that affects female reproductive success via interactions with both mutualists and antagonists. We discuss the interplay between the resource status and selection imposed on female reproductive success and its likely role in the evolution of sex-allocation strategies, especially andromonoecy.

## INTRODUCTION

A central goal of plant evolutionary biology is to understand how selection acts on phenotypic traits in wild populations. The functional significance of variation in floral traits has attracted particularly keen attention (Caruso et al., 2019), not least because they vary so strikingly among species, populations, and individuals, but also because they can often be linked directly to key components of reproductive success and thus fitness (Gómez and Zamora, 2000; Maad, 2000; Ågren et al., 2013; Sletvold et al., 2013). For instance, the rate at which ovules are fertilized by self-versus outcross pollen (which affects the female component of reproductive success) may depend on factors such as the attractiveness of inflorescences and flowers (Waser, 1983; Caruso et al., 2019) and the manner in which flowers manipulate pollinator behavior (Schiestl and Johnson, 2013). Similarly, the ability of plants to disperse their pollen effectively to other flowers in the population (affecting male reproductive success) also depends on inflorescence and floral traits (Conner et al., 1996; Hodgins and Barrett, 2008). While interactions with mutualists thus likely play an important role in shaping floral evolution, floral traits may also influence fitness via their antagonistic interactions with herbivores, florivores, and seed predators, through both their male and female components of fitness (Strauss and Whittall, 2006). For instance, both pollinators and antagonists have been shown to impose selection in the same or in different directions on floral color (Frey, 2007; Carlson and Holsinger, 2010; Ehrlén et al., 2012), floral size (Gómez, 2003; Pérez-Barrales et al., 2013), floral scent (Schiestl et al., 2011; Knauer and Schiestl, 2017), and nectar production (Kessler et al., 2015; Parachnowitsch et al., 2019). However, much less is known about how selection operates on traits that could be regarded as ancillary to flowering and floral function.

The height of the vegetative stalk on which the flowers and fruits develop is an ancillary trait that might affect plant fitness in several ways, positively or negatively (reviewed in Harder and Prusinkiewicz, 2013). Positive selection, through male fitness favoring longer branches or inflorescence stalks, has been shown for wind-pollinated species (Tonnabel et al., 2019), and a positive correlation between stalk height and pollinator visitation rate, which may strongly affect siring success, has been found in many animal-pollinated species (Galen, 1989; O’Connell and Johnston, 1998; Gómez, 2003; Sletvold and Ågren, 2015; Diniz et al., 2019). In species with wind-dispersed seeds, floral stalk height can also have a strong positive effect on seed-dispersal distances. For example, Greene and Johnson (1989) and Soons et al. (2004) showed that horizontal wind velocity and seed release height (i.e., stalk height) both affected seed dispersal. Stalk height may also negatively affect a plant’s female component of fitness if flowers on tall stalks are more easily seen or accessed by herbivores and seed predators, as has been found, for example, in *Castilleja linariaefolia* (Cariveau et al., 2004), *Erysimum mediohispanicum* (Gómez, 2003), and *Primula farinose* (Ågren et al., 2013).

Selection on floral stalk height via its effect on both pollen and seed dispersers and potential antagonists also needs to be considered in the context of the marginal costs of stalk production under different resource levels, and in different vegetation contexts. First, to the extent that stalk height affects fitness positively, we should expect larger plants with more resources to produce taller stalks. In contrast, plants that have lost carbon and nutrients to prior herbivory, for instance, may be constrained to produce shorter stalks (Waite and Hutchings, 1982; Weiner, 2004), with potentially negative consequences for pollen and/or seed dispersal (Donohue, 1999; Sletvold et al., 2013). Second, as with overall plant height itself, the fitness implications of stalk height are likely to depend on the height of the surrounding vegetation, with plants producing taller stalks to maintain height above neighboring plants (Sletvold et al., 2013). Thus, while stalk height is an apparently simple quantitative trait, its expression is likely to be the outcome of responses to selection on norms of reaction to a plant’s resource status (e.g., its overall size and history of herbivory), the height and density of vegetation in which it is expressed, and the extent to which different flowers vary in their sex allocation.

Optimal stalk height may differ for flowers with different sex allocation (i.e., the proportion of their resources committed to seed versus pollen production) if greater height benefits fitness via interactions with mutualists and antagonists more through one sex than the other. In many species, the sex allocation varies considerably at both the individual and inflorescence or floral levels, and we might expect stalk height and sex allocation to covary in such species. For instance, Pickup and Barrett (2012) found that the height of stalks of males of wind-pollinated dioecious *Rumex hastatulus* was greater than that of females during flowering, but that female stalk height was greater during fruiting and seed dispersal. To our knowledge, however, the differential selection of stalk height as a function of sex allocation has hitherto not been investigated in any hermaphroditic or monoecious species. This represents an important gap. Sex allocation is expected to vary as a function of plant size in many hermaphroditic species (de Jong and Klinkhamer, 1989; Klinkhamer et al., 1997), not only because larger plants have more resources and can thus potentially allocate more to the more costly sex (a so-called ‘budget effect’ of size), but also because taller plants are better able to disperse seeds and/or pollen (a ‘direct effect’ of plant size) (Klinkhamer et al., 1997).

Here, we explore the effects of floral stalk height on female components of plant reproductive success in the insect-pollinated and wind-dispersed alpine perennial herb *Pulsatilla alpina* (Ranunculaceae). This species displays wide variation among individuals not only in floral stalk height but also in floral sex allocation (the number of stamens and pistils per flower), plant size (and thus resource status), and the impact on resource status by herbivores. Flowers of *P. alpina* are presented to pollinators early in the growing season, as soon as snows melt. They are pollinated largely by generalist dipteran pollinators (Appendix S1, see Supplementary Data), which might be attracted to flowers on taller stalks. However, they are also visited by a specialist dipteran seed predator that lays its eggs in the gynoecium and whose larvae eat the seeds (technically the achenes), potentially mitigating against the benefits of greater height during flowering. In addition, stalk height is likely to have a positive effect on seed dispersal by wind, particularly when plants are growing in tall vegetation.

We first describe variation in floral stalk height and other floral characters within and among populations of *P. alpina* in the French-Swiss Alps. We then assess phenotypic selection on the female components of reproductive success in terms of morphological variation in flowers and floral stalks. In particular, we address the following questions. (1) How does stalk height vary among populations, and, in particular, as a function of vegetation height? We expected plants in taller vegetation to have taller stalks. (2) Within a population, how does stalk height vary across developmental stages, and how does its development correspond to that of other floral traits? To the extent that taller stalks during fruiting and seed dispersal have a positive influence on plant fitness, we expected stalk height to increase between flowering and fruiting in this hermaphroditic species, as found previously for females of a dioecious herb (Pickup and Barrett, 2012). (3) Does stalk height covary with floral sex allocation? A finding of greater stalk height for flowers with greater relative allocation to one sex would be consistent with fitness through that sex benefitting more from floral height than through the other. (4) To what extent might herbivory on leaves and vegetative shoots impact the height of floral stalks plants are able to produce? If producing tall stalks is costly, we expected simulated herbivory to cause plants to produce shorter stalks. And (5) How does stalk height influence both the ovule fertilization rate and the rate of seed predation? If both of these rates are greater in flowers on taller stalks, this would provide evidence for different components of selection via pollinators and seed predators operating in different directions.

## MATERIAL AND METHODS

### Study species

*Pulsatilla alpina* (L.) Delarbre is a perennial hemicryptophyte distributed from sub-alpine to alpine grassland in central Europe (Lauber et al., 2018). Several shoots emerge from a perennial rhizome soon after the snowmelt, from early May to July; these shoots may be vegetative or reproductive. The plants vary greatly in both above-ground size (number of leaves and flowers) and in the size of the persistent underground rhizome. Above-ground herbivory in the study populations is mainly the result of direct consumption or trampling by cattle, usually in late summer.

A single flower with white showy tepals is produced at the apex of its own floral shoot. Individuals produce both male and hermaphroditic flowers (the species is thus ‘andromonoecious’), with wide variation also in their number of pistils and stamens per flower, i.e., hermaphrodite flowers vary quantitatively in their sex allocation. Phenotypically male flowers bear no pistil, and the pistil number in hermaphroditic flowers varies from about ten to about 400. Each pistil contains only one ovule. The stamen number varies from about 150 to about 400 in both male and hermaphroditic flowers. The flower number varies from one to 20 flowers among individuals and populations. Fruits ripen to produce achenes with an elongated pappus that promotes dispersal by wind (Muller-Schneider, 1986; Vittoz and Engler, 2007). After achenes are dispersed in autumn, the above-ground parts of the plants wither, but individuals persist underground as a rhizome until the next spring.

Flowers of *P. alpina* are visited by both dipteral pollinators and seed predators. The main pollinators of *P. alpina* are house flies and syrphids (Appendix S1), which visit the flowers for pollen (Szentpéteri et al., 2008). Flowers of *P. alpina* are also visited by a monophagous dipteran seed predator, *Phytomyza* sp.. These flies mate in the flower, the female adults oviposit on the pistils, and the larvae predate the pollinated ovules during the fruiting stage, as has been found for other *Phytomyza* species that predate seeds within the achenes (Winkler et al., 2009).

### Survey on variation in stalk height among populations

We conducted our survey and experiments in 13 populations in the pre-Alps of the Canton Vaud, Switzerland, over three consecutive years from 2019 to 2021 (see Appendix S2 in Supplementary Data for the details of each population). To characterize variation in stalk height among populations, we sampled eleven populations at the end of the growing season. In each population, we established four to five transects across the population and permanently marked them. Within each one-meter-wide transect, we sampled the highest floral stalk at the fruiting stage from around 20 individuals of *P. alpina*, i.e., we sampled a total of around 80 to 100 individuals per population, though in populations with a high proportion of non-flowering individuals or low population density (i.e., populations S1+, S4, and LS2), only around 50 flowering individuals were sampled. In all, we sampled stalks from 757 individuals from 41 transects at the end of the growing season. We measured the stalk height as the distance between the bottom of the stalk aboveground and the receptacle of the flower to which the achenes are attached.

To determine the effect of intrinsic and extrinsic factors on the variation of stalk height among populations, we recorded the total flower number of sampled individuals and vegetation height along the transects in the early and late growing season. We used the total flower number to infer a plant’s resource status (larger plants produce more flowers). We measured the vegetation height in a semi-quantitative way by assigning the vegetation along the transects into six height categories, i.e., > 100 cm, 61 to 100 cm, 31 to 60 cm, 11 to 30 cm, 1 to 10 cm, and zero cm, and recorded the cover of each height class on a six-interval scale: < 1%, 1% to 5%, 5% to 25%, 25% to 50%, 50% to 75%, and 75% to 100%. We calculated the mean height for each transect by weighting height classes in terms of the centroid of their cover class for the transect.

### Variation in stalk height through developmental stages, and correlation with other floral traits at the flower level

To characterize variation in stalk height at the flower level, we followed the development of floral stalks in 60 flowers from 20 individuals that varied in plant size and flower number (from one to 13 flowers) through the course of the growing season in population S1. We recorded each floral stalk every three to five days, from the budding stage toward the end of the flowering stage, and every seven to 20 days until the end of the fruiting stage. We defined the period of the budding stage as beginning with the emergence of the floral stalk and ending when the flower opened its tepals. The flowering stage was defined as beginning when flowers opened and ending when they wilted. The fruiting stage began as flowers wilted and ended with the commencement of fruit dispersal.

To evaluate the correlation of flower height with other important floral traits of the species, we recorded tepal length, pistil number, and stamen number at the end of the flowering stage. We measured the length of the largest tepal of the flowers, which is at the outer-whorl in most of the cases, from the tip of the tepal toward its bottom attached to the receptacle. We conducted measurements on dry days when tepals were fully expanded. To measure sex allocation of each flower because the species possesses a great variation in the number of pistils and stamens within a flower. We photographed the flowers and counted the pistil number from top-viewed photos and the stamen number from side-viewed photos then multiplied by two.

### The effect of leaf removal on floral stalk height

To manipulate the potential resource availability of individuals for investment towards reproduction, we conducted a leaf-removal experiment in two populations (LL3, 2019, N = 101; and S1, 2020, N = 72) in which herbivory was deemed to be low, based on personal observations in 2018. Around 12 flowering individuals bearing one, two, or > two flowers were arbitrarily chosen and labeled with a metal tag on the ground at the beginning of the flowering season within each subplot in each population and were randomly assigned to either a leaf-removal (LR) or a control (C) treatment, by a random draw of lots. For the LR treatment, two-thirds of the total leaves were removed at the beginning of the flowering season. A second round of leaf-removal was performed a month later to ensure that all the newly emerged leaf stalks were defoliated. C plants were not damaged. Each flower was labeled with a tag and photographed. We measured the height of all the floral stalks of the individuals at the end of the flowering stage with the same method described in the previous section.

### Selection differentials and selection gradients on stalk height for female fitness components before seed dispersal

To evaluate the female component of selection on floral stalk height, we quantified achene fate for 322 seed families collected in four populations in 2019, i.e., population S1 (N = 95 flowers from 45 individuals), population S2 (N = 74 flowers from 42 individuals), population LL1 (N = 62 flowers from 42 individuals), and population LL4 (N = 91 flowers from 41 individuals). We marked the flowers with a tag at the beginning of the flowering season, chosen arbitrarily from different individuals varying in their size. We measured the same floral traits at the end of the flowering stage, as described in the previous section, i.e., stalk height, tepal length, and the number of stamens. We calculated the number of pistils as the sum of the number in each of the three achene categories in each flower (see below for details). We collected the achenes of the labeled flowers at the end of the growing season before they started to disperse. Because male flowers bore no achene, we used only hermaphroditic flowers in the analysis.

To quantify the pre-dispersal components of female reproductive success, we separated achenes into three categories: unfertilized, predated, and mature, based on morphological assessment (see Appendix S3 in the Supplementary Data for detailed description). We interpreted the seed fertilization rate as a reflection of pollinator visitation, autonomous selfing, and resource limitation (see Discussion). The seed predation rate reflects selection imposed mainly by the *Phytomyza* seed predator, but may also be affected by selection via organisms at higher trophic levels, e.g., parasitoid wasps, which parasitize the seed predators and thus potentially reduce the seed predation rate. The seed maturation rate describes the proportion of all achenes that were fertilized and not predated. Lastly, we used mature achene number as one of the fitness components reflecting the contribution to the gene pool of the next generation via the female function. In summary, we considered four fitness components in our analyses: the number of mature achenes; the fertilization rate (the sum of predated and mature achenes divided by all achenes produced in the flower); the achene predation rate (the number of predated achenes divided by the number of fertilized achenes); and achene maturation rate (the number of mature achenes divided by all achenes produced in the flower). For greater clarity in interpreting directions of selection, we refer to the rate of non-predation (calculated as 1 – predation rate) instead of the predation rate in what follows.

### Statistical analysis

We conducted all analyses within the R statistical framework v 4.0.3 (R Core Team, 2021), using package lme4 for (generalized) linear mixed models (Bates et al., 2015). We evaluated the fit of each model with the R package DHARMa (Hartig, 2019).

We used a linear mixed model to evaluate the dependence of stalk height on flower number and vegetation height during early and late season, with population, flower number, and average vegetation height during early and late season as fixed effects. We considered transects as a random effect that grouped different individuals. To evaluate the relationship among stalk height, tepal length, pistil number, and stamen number, we calculated the Pearson correlation for each trait pair. To evaluate the effect of leaf-removal on stalk height in the next season, we used a linear mixed model, with population, treatment, and their interaction as fixed effects. We considered individual as a random effect to account for the fact that more than one stalk was sampled from some individuals.

To evaluate the phenotypic selection differentials and selection gradients on stalk height for the four components of female reproductive success, we used generalized linear mixed models for fertilization rate, rate of non-predation, and seed maturation rate and a linear mixed model for relative female fitness. The difference between a selection differential and a selection gradient is that the former measures the total strength of selection on the trait (direct and indirect effect via correlation with other traits), while the latter measures only the strength of direct selection (partial effect) on the trait (Brodie et al., 1995). We calculated the relative mature achene number by dividing the number of mature achenes of each flower by the mean number of mature achenes across the population. We standardized stalk height, tepal length, pistil number, and stamen number for each population to a mean of zero and a standard deviation of one.

To determine the linear (S) and quadratic (Cii) selection differentials, which are the total strength of selection on the trait (direct and indirect), we set linear and quadratic terms for stalk height as fixed effects and considered the interactions with the population in each simple regression model (Lande and Arnold, 1983; Matsumura et al., 2012). We used generalized linear mixed models with a logistic function for binomial response variables, i.e., fertilization rate, rate of non-predation, and seed maturation rate, and a linear mixed model for relative mature achene number. We used the ‘emtrends’ function in the R package emmeans to extract the regression coefficients and standard errors of the general effects and each population from the models (Lenth, 2020). Regression coefficients and their standard errors from the generalized models of each population were adjusted by multiplying the value with a constant to approximate the selection differentials (Janzen and Stern, 1998). The constant is the average of W(z)[1 - W(z)], where W(z) is the predicted fitness value for each individual in the studied population using the estimated logistic regression coefficients. A significant linear differential indicates a directional selection on the trait, and a significant quadratic differential in conjunction with a local maximum and minimum indicates stabilizing or disruptive selection, respectively. A significant interaction between a floral trait and the population indicates a difference in selection among populations. For all quadratic differentials, we multiplied the regression coefficient by two to obtain the correct estimate of stabilizing or disruptive selection (Stinchcombe et al., 2008). We set individual identity as a random effect to account for the fact that more than one flower was sampled in some individuals in all the regression models. In addition, we set flower identity as a random effect in the generalized models to account for the fact that the unit of the binomial response variables (e.g., fertilization rate) is one achene grouped with the other achenes of the same flower.

To determine the linear (*β*) and quadratic (*γ*_*ii*_) selection gradients, we set linear and quadratic terms of the four floral traits as fixed effects and considered the interactions with population in each multiple regression model (Lande and Arnold, 1983; Morrissey, 2014). This allowed us to extract the partial effect on the fitness components by each trait. We used generalized linear mixed models with a logistic function for binomial fitness components and a linear mixed model for relative mature achene number. We set random effects as described above for the simple regression models. Regression coefficients and their standard error from the generalized models were adjusted by multiplying the value with a constant, as described above (Janzen and Stern, 1998). For all quadratic gradients, we multiplied the regression coefficients by two to obtain the correct estimate of stabilizing or disruptive selection (Stinchcombe et al., 2008).

## RESULTS

### Among-population variation in floral stalk height

Across all individuals sampled, floral stalk height varied from 21 to 72 cm (mean and standard deviation: 46.5 ± 8.71), with substantial and significant (P < 0.001) variation among populations, from a minimum in population S4 (34.9 ± 6.84 cm) to a maximum in population LM1 (52.9 ± 9.28 cm) (Fig. 1A). Individuals producing more flowers also produced taller stalks (P < 0.001) (Fig. 1B). Individuals in taller vegetation at late-season produced taller stalks (P < 0.01) (Fig. 1C), but not at early season (Appendix S4, see Supplementary Data). These patterns were consistent within and among populations (Fig. 1B and 1C).

**FIGURE 1.**
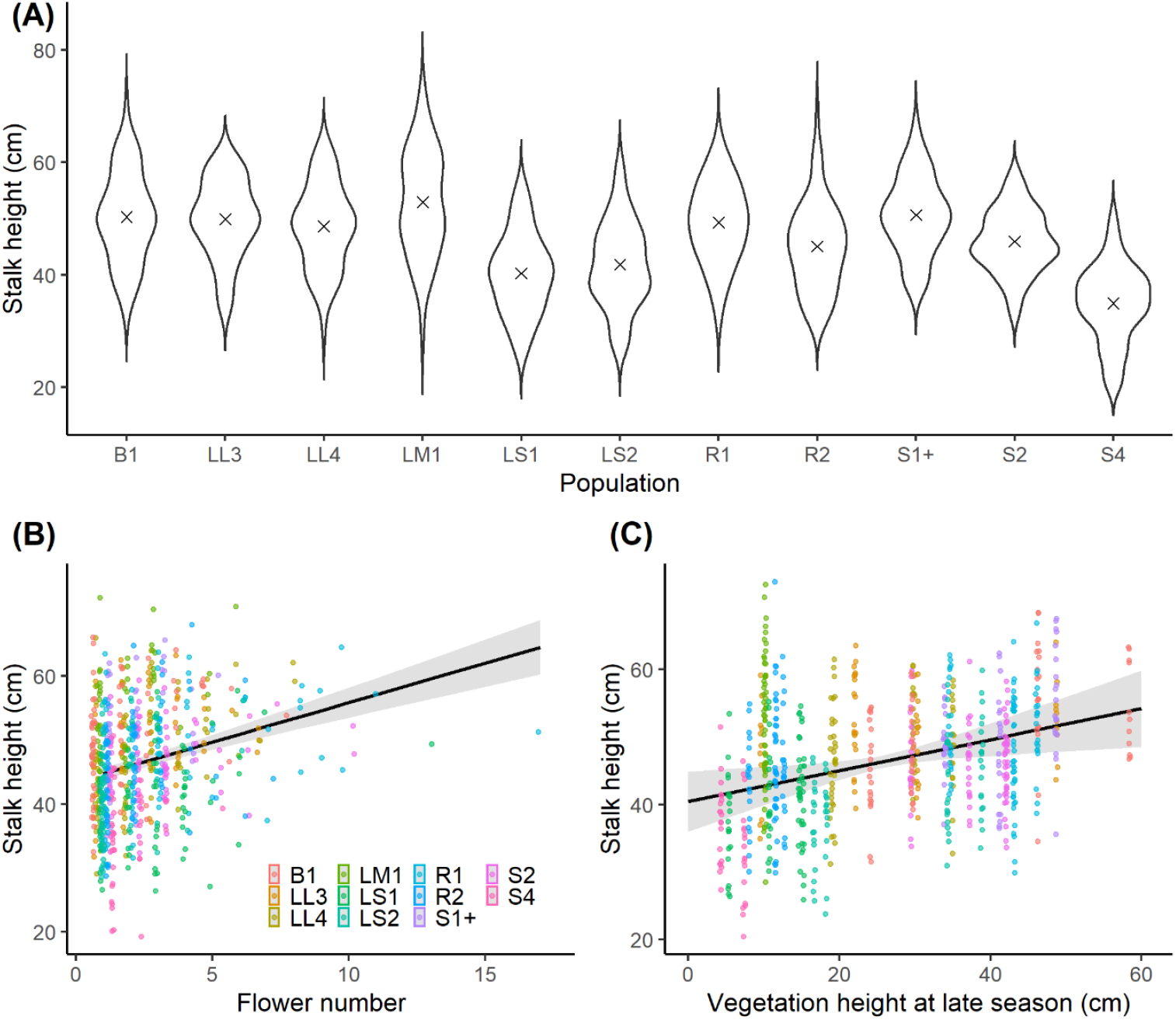
(A) Violin plot showing the variation in stalk height at the end of the growing season across eleven populations studied. The mean of each population is indicated by crosses. (B) Predicted values of stalk height as a function of flower number. (C) Predicted values of stalk height as a function of late-season vegetation height. Data points from different populations were shown in different colors in (B) and (C). The shaded ribbon indicates the 95% confidence interval of the regression lines in (B) and (C).

### Variation in floral stalk height over time, and its correlation with other floral traits

We fitted a local polynomial regression on stalk height through developmental stages to capture the dynamics of stalk height (Fig. 2A). Stalks continued to elongate during the budding and flowering stage, roughly the first 20 days after their emergence aboveground, but showed no further elongation after the end of flowering. Stalk height correlated positively with pistil number (r = 0.65, P < 0.001) and tepal length (r = 0.51, P < 0.001) (Fig. 2A and 2B). The stalk height for male and hermaphroditic flowers was 30.4 ± 6.69 and 49.9 ± 6.96 cm, respectively, at the end of the male stage, and was 32.8 ± 6.18 and 55.8 ± 10.2 cm, respectively, at the end of the fruiting stage.

**FIGURE 2.**
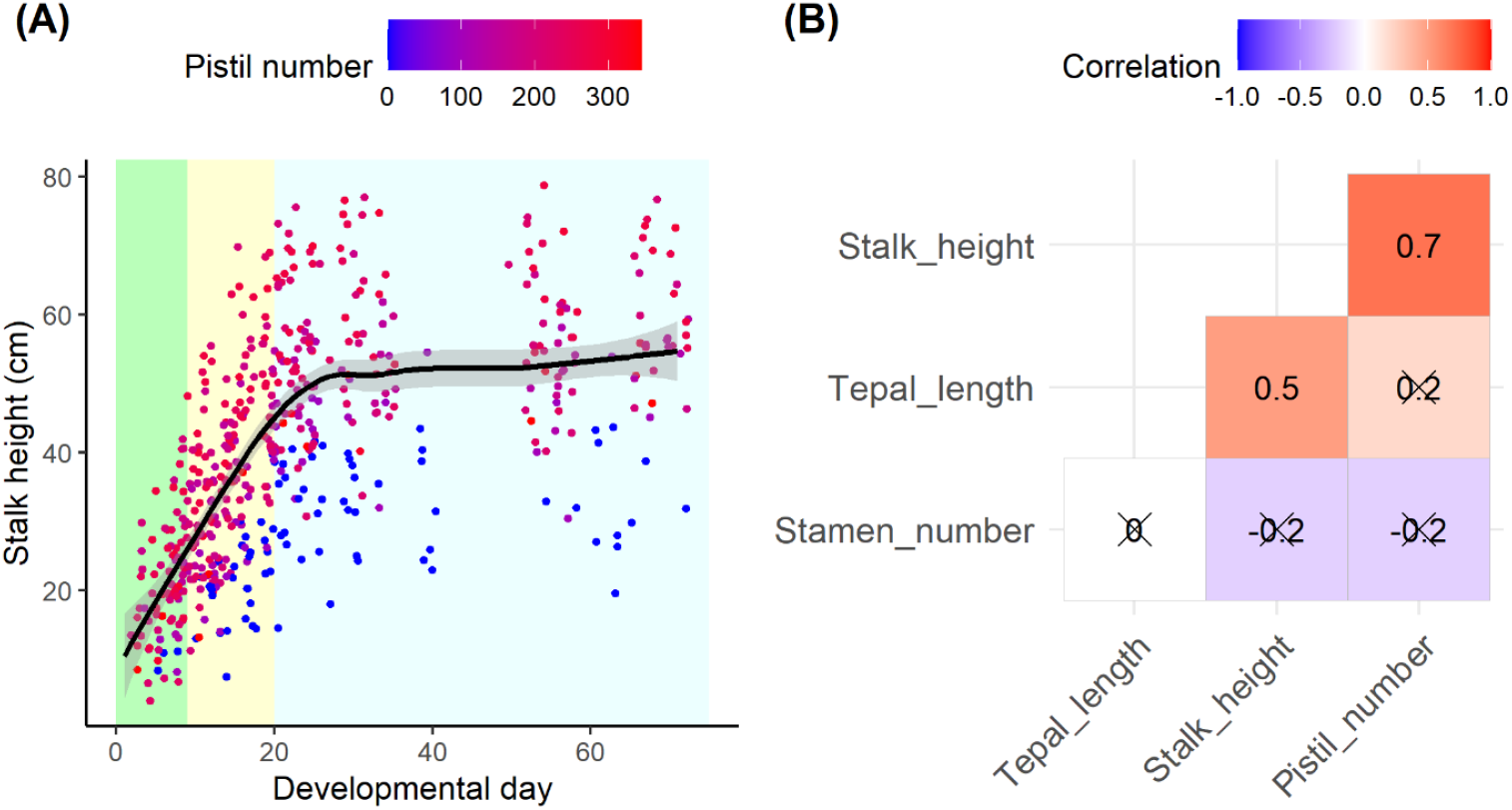
(A) Dynamics of the height of 60 floral stalks across the season from bud stage to fruiting stage. The color gradient of the points indicates the number of pistils of the flower. Bud stage, flowering stage, and fruiting stage are shown in light green, yellow, and blue respectively. The shaded ribbon indicates the 95% confidence interval of the smoothed lines. (B) Correlation matrix for stalk height, tepal length, pistil number, and stamen number. Floral traits were measured at the end of the flowering stage. Non-significant correlations (P > 0.05) are shown under a cross.

### Effects of leaf removal on floral stalk height

Plants subject to leaf-removal produced significantly shorter stalks the following season in two populations than did control plants (33.7 ± 9.04 and 39.4 ± 12.4 cm in LR and C treatments, respectively; P < 0.01). The interaction between population and treatment was not significant (Fig. 3 and Appendix S5 in the Supplementary Data).

**FIGURE 3.**
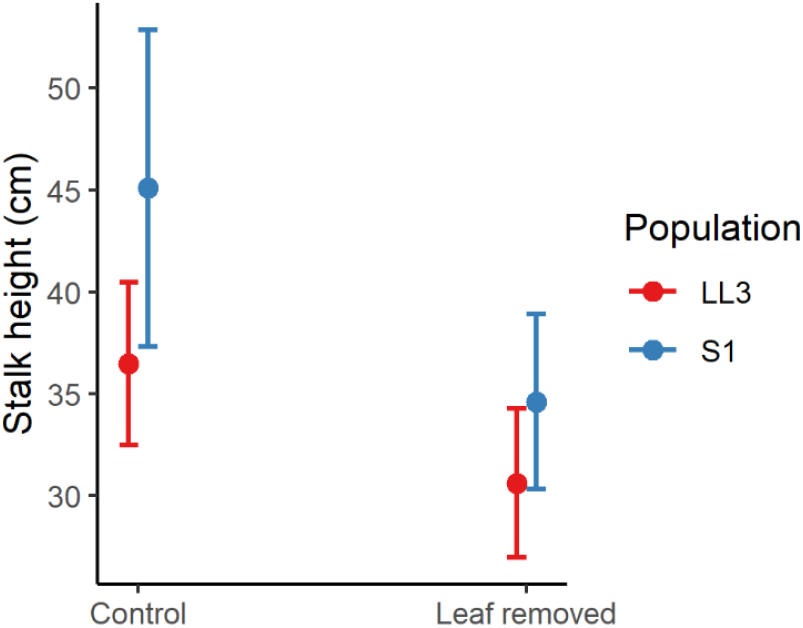
Predicted values of stalk height in year *t* after leaf removal (or not) in year *t* – 1. Results are shown for two populations (see inset legend). The upper and lower bar indicates the 95% confidence interval. Plants from the leaf removal treatment produced shorter stalks in the next season in both the two populations (P < 0.01).

### Selection differential on floral stalk height for pre-dispersal components of female reproductive success

In total, 67,798 achenes were counted to determine the rate of fertilization, the rate of non-predation, the rate of seed maturation, and the number of mature seeds. The mean and standard deviation of the fitness components and stalk height of the four populations can be found in Appendix S6 (see Supplementary Data).

Overall, our models found largely consistent trends for the selection differentials on stalk height across the studied populations in the four fitness components (Table 1, Fig. 4). There was a tendency towards positive quadratic effects of stalk height on fertilization rate (general linear coefficient = 0.0722, P > 0.05; quadratic coefficient = 0.109, P < 0.01), rate of non-predation (general linear coefficient = -0.121, P > 0.05; quadratic coefficient = 0.119, P < 0.05), and seed maturation rate (general linear coefficient = 0.006, P > 0.05; quadratic coefficient = 0.133, P < 0.01). There was a tendency towards a positive linear effect of stalk height on relative mature seed number across the four populations (general linear coefficient = 0.186, P < 0.001; quadratic: γ = 0.004, P > 0.05) (see Table 1 and Fig. 4 for the values and illustrations of the selection differentials of each population, and Appendix S7 (see Supplementary Data) for figures showing the general effects). The interaction with population was non-significant for both linear and quadratic terms for all the fitness components, with a marginally non-significant interaction between population and the quadratic terms for the rate of non-predation (Fig. 4B, P = 0.064). There was a single minimum value for the fertilization and seed maturation rates within the phenotypic range of stalk height in each of the four populations predicted by the models, indicating a largely consistent pattern of a disruptive selection on stalk height for the two fitness components (Fig. 4). For the rate of non-predation, a single minimum value was found in three populations and a maximum was found in one population (Fig. 4), pointing to disruptive selection in the former three and stabilizing selection in the latter. The pattern of disruptive selection on stalk height in the fertilization rate, the rate of non-predation, and the seed maturation rate may have been driven to some extent by the influence of extreme phenotypes, as evidence for disruptive selection was substantially weakened when extreme phenotypes were removed from the analysis (see Appendix S8 in the Supplementary Data). Lastly, a maximum value of relative mature seed number was found at the positive end of the range of standardized stalk height in three populations, pointing to positive directional selection (Fig. 4).

**TABLE 1.** Selection differentials and selection gradients on stalk height of four fitness components in four studied populations. Linear (S) and quadratic (Cii) selection differentials of stalk height and the standard errors were derived from single regression models. Linear (β) and quadratic (γii) selection gradients and the standard errors of four floral traits were derived from multiple regression models with the other floral traits. Significant values (P < 0.05) are in bold. Marginally non-significant values (0.05 < P < 0.1) are in italic.

|  | Fertilization rate |  |  |  | Rate of non-predation |  |  |  | Seed maturation rate |  |  |  | Relative mature seed number |  |  |  |
| --- | --- | --- | --- | --- | --- | --- | --- | --- | --- | --- | --- | --- | --- | --- | --- | --- |
|  | LL1 | LL4 | S1 | S2 | LL1 | LL4 | S1 | S2 | LL1 | LL4 | S1 | S2 | LL1 | LL4 | S1 | S2 |
| S<br>(Linear selection differential) | 0.012<br>$\pm 0.023$ | 0.014<br>$\pm 0.023$ | 0.027<br>$\pm 0.024$ | 0.009<br>$\pm 0.027$ | 0.008<br>$\pm 0.024$ | -0.04<br>$\pm 0.026$ | -0.033<br>$\pm 0.028$ | -0.034<br>$\pm 0.03$ | 0.008<br>$\pm 0.022$ | -0.009<br>$\pm 0.023$ | 0.007<br>$\pm 0.026$ | -0.002<br>$\pm 0.026$ | *0.147*<br>$\pm 0.078$ | *0.135*<br>$\pm 0.073$ | **0.311**<br>$\pm 0.074$ | *0.152*<br>$\pm 0.085$ |
| C<sub>ii</sub><br>(Quadratic selection differential) | *0.073*<br>$\pm 0.039$ | 0.017<br>$\pm 0.026$ | 0.024<br>$\pm 0.028$ | *0.071*<br>$\pm 0.038$ | **0.115**<br>$\pm 0.041$ | *0.059*<br>$\pm 0.032$ | 0.026<br>$\pm 0.032$ | -0.029<br>$\pm 0.04$ | **0.101**<br>$\pm 0.038$ | 0.034<br>$\pm 0.026$ | 0.034<br>$\pm 0.03$ | 0.025<br>$\pm 0.035$ | 0.175<br>$\pm 0.133$ | 0.036<br>$\pm 0.081$ | -0.114<br>$\pm 0.087$ | -0.063<br>$\pm 0.105$ |
| $\beta$<br>(Linear selection gradient) | -0.004<br>$\pm 0.024$ | 0.009<br>$\pm 0.024$ | 0.011<br>$\pm 0.026$ | 0.011<br>$\pm 0.027$ | **0.048**<br>$\pm 0.024$ | -0.041<br>$\pm 0.027$ | -0.018<br>$\pm 0.028$ | 0.007<br>$\pm 0.032$ | 0.014<br>$\pm 0.023$ | -0.015<br>$\pm 0.024$ | 0<br>$\pm 0.028$ | 0.02<br>$\pm 0.029$ | 0.007<br>$\pm 0.073$ | -0.038<br>$\pm 0.071$ | 0.095<br>$\pm 0.077$ | -0.023<br>$\pm 0.087$ |
| $\gamma_{ii}$<br>(Quadratic selection gradient) | *0.067*<br>$\pm 0.039$ | 0.01<br>$\pm 0.025$ | 0.02<br>$\pm 0.03$ | **0.021**<br>$\pm 0.03$ | **0.086**<br>$\pm 0.038$ | *0.054*<br>$\pm 0.029$ | 0.018<br>$\pm 0.031$ | -0.041<br>$\pm 0.038$ | **0.087**<br>$\pm 0.037$ | 0.027<br>$\pm 0.025$ | 0.027<br>$\pm 0.031$ | 0.027<br>$\pm 0.034$ | *0.214*<br>$\pm 0.12$ | 0.023<br>$\pm 0.072$ | -0.03<br>$\pm 0.084$ | 0.031<br>$\pm 0.097$ |

**FIGURE 4.**
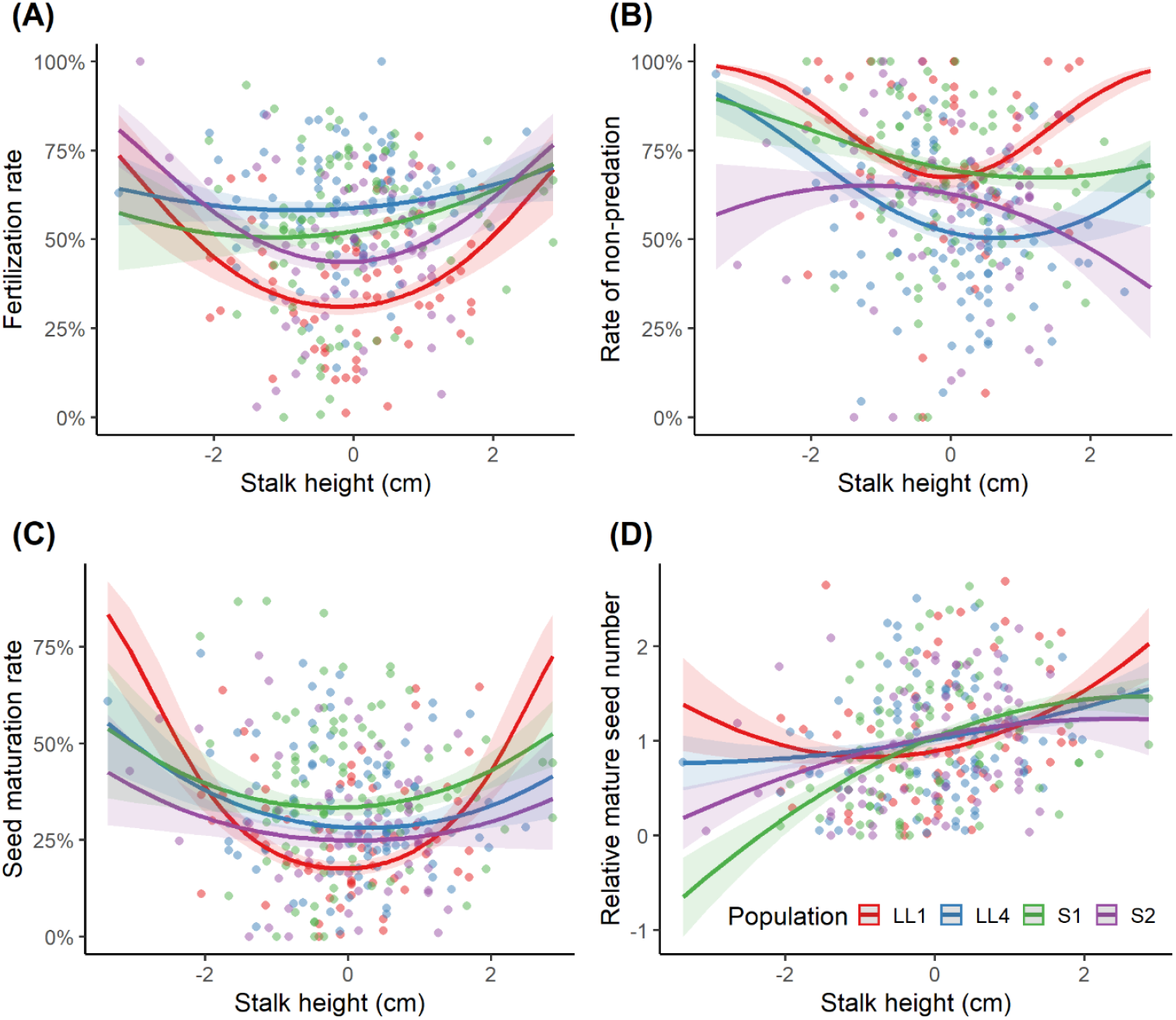
Visualization of the relationship between standardized stalk height at the end of the flowering stage and (A) fertilization rate, (B) rate of non-predation, (C) seed maturation rate, and (D) relative mature seed number among populations from single regression models evaluating the selection differentials. Raw data points and regression lines of the populations studied are shown in different colors (interactions between stalk height and population were all non-significant). The shaded ribbons indicate the standard errors of the regression lines. See Table 1 for the significant values of the linear and quadratic selection differentials.

### Selection gradients on floral stalk height for pre-dispersal components of female reproductive success

We focused on the main results of the selection gradient on floral stalk height in the four female fitness components from the multiple regression models. A complete table of the results of the multiple regression models for all traits can be found in Appendix S9 (see Supplementary Data).

Overall, the selection gradients on stalk height were largely consistent across the studied populations for the four fitness components, as also found for the selection differentials (Table 1). There was a tendency towards positive quadratic effects for the regression of stalk height on fertilization rate (linear coefficient = 0.045, P > 0.05; quadratic coefficient = 0.102, P < 0.05), rate of non-predation (linear coefficient = 0.01, P > 0.05; quadratic coefficient = 0.083, P < 0.05), and seed maturation rate (linear coefficient = 0.028, P > 0.05; quadratic coefficient = 0.115, P < 0.01). Neither linear nor quadratic terms were significant for regressions of stalk height on relative mature seed number (linear coefficient = 0.01, P > 0.05; quadratic coefficient = 0.03, P > 0.05) (Table 1 and Appendix S9). The interaction with population was non-significant in both linear and quadratic terms for fertilization rate, seed maturation rate, and relative female fitness, while the interaction with population fell just short of significance for the rate of non-predation (P = 0.081 and 0.082 for linear and quadratic, respectively) (Appendix S9).

The direction and shape of selection differentials and selection gradients of each population were largely consistent, though the pattern was statistically significant for only some of them. This may be because of a variation in the strength of selection, or due to low statistical power. Nonetheless, the consistency of the effects of stalk height on the four fitness components among populations was reflected in highly significant values for the general pattern (Appendix S7 and S9).

## DISCUSSION

Our study has revealed evidence for unusual disruptive selection on floral stalk height in *P. alpina* in terms of three different components of reproductive success. Our results also indicate that floral stalks are costly to produce and that investment in such stalks should likely be attributed largely to female rather than male components of reproductive success, a feature that may help to explain the andromonoecious sexual system of this perennial herb.

### Disruptive selection on stalk height for three components of female reproductive success

Our results point to disruptive selection on floral stalk height in *P. alpina*, with flowers on taller and shorter stalks having a higher fertilization rate, lower seed predation rate, and higher seed maturation rate than those of intermediate height. It may be significant that we found a positive selection differential but not a positive selection gradient for relative mature seed number across the populations sampled (Table 1, Fig. 4, and also Appendix S7, S9). Given that the selection differential measures the total strength of selection on stalk height (direct and indirect effect via correlation with other traits), and the selection gradient measures only the strength of direct selection (partial effect) on stalk height (Brodie et al., 1995), it is possible that the higher relative mature seed number associated with taller stalks (i.e., positive selection differential) may be an indirect effect through a positive correlation with, for instance, pistil number.

The disruptive selection on stalk height in terms of fertilization rate (Table 1, Fig. 4A and Appendix S7A) may be a result of both pollinator preference and facilitated or autonomous selfing. On the one hand, several previous studies have found a positive selection on stalk height by the pollinators on female fitness (Galen, 1989; O’Connell and Johnston, 1998; Sletvold et al., 2015), and a strong preference for flowers on higher stalks in the grassland plant community has been found in several syrphid fly species, which are the main pollinators of *P. alpina* (Klecka et al., 2018). On the other hand, *P. alpina* individuals produce actinomorphic flowers with pistils surrounded by numerous stamens, as is found in many basal eudicots. The pistils in the outer whorl that are directly beside the stamens may be subject to self-pollination, either autonomously or facilitated by pollinators. As short stalks usually have fewer pistils, it is possible that the high fertilization rate in short stalks is a consequence of such selfing. If so, these two mechanisms may ultimately lead to the observed disruptive selection in the fertilization rate, with the degree of inbreeding depression determining the direction of the selection (e.g., Briscoe Runquist et al., 2017). Estimates of the selfing rate in flowers with different allocation strategies and stalk heights would be worthwhile to test this hypothesis.

On the other hand, our finding of disruptive selection on stalk height due to seed predation differs from that of most studies, which have tended to reveal directional negative selection on stalk height by antagonists, from invertebrates to ungulates (Gómez, 2003, 2008; Cariveau et al., 2004; Ehrlén et al., 2012; Ågren et al., 2013). However, these other studies have typically considered only directional selection in their analyses, i.e., only linear terms were included in the relevant statistical models analyzed and disruptive or stabilizing selection was not formally tested (e.g., Cariveau et al., 2004; Ehrlén et al., 2012). In one of the few studies that did consider quadratic terms in the multiple regression, Gomez (2008) found evidence for stabilizing selection on stalk height in the rate of seed predation in *Erysimum mediohispanicum*, in contrast to our finding of disruptive selection. Furthermore, it is likely that the seed predation rate in *P. alpina* depends not only on the preference of seed predators but also on that of the predators or parasites such as parasitoids of the seed predators (personal observations), as in several other systems (Molau et al., 1989; Gómez and Zamora, 1994). However, the host-feeding strategies of parasitoids are complicated, and most studies have focused only on olfactory cues (Jervis and Kidd, 1986; Giunti et al., 2015). Thus, how parasitoids play the role in the selection of a trait such as stalk height remains largely unexplored.

Whatever its causes, a striking feature of the patterns pointing to disruptive selection on floral stalk height is their consistency among populations, not only through pollinators but also through seed predators. Some studies have reported counteracting selection by pollinators and seed predators (Gómez, 2007; Ehrlén et al., 2012; Thomann et al., 2018), while others have found that the floral trait under investigation was under selection by one of the agents (Cariveau et al., 2004). Furthermore, some studies have shown a geographic mosaic (*sensu* Thompson, 1999) in the direction and shape of selection imposed by different agents such as pollinators, herbivores, or seed predators (Gómez and Zamora, 2000; Gómez et al., 2009; Ågren et al., 2013). In contrast, we found a rather constant pattern across the four populations studied. It seems likely that stalk height in *P. alpina* varies not so much because of differences among populations in the strength and direction of selection due to mutualists or antagonists, but rather as a result of heterogeneity in, for example, the sex allocation of their flowers, competition with other plants in the surrounding vegetation, the resource status of the plants, and perhaps differences in selection between male and female components of fitness.

### Female sex allocation, inferred cost of stalk height, and the evolution of the sexual system

Our results suggest that stalk height affects the female component of reproductive success in *P. alpina* and may be less relevant to male reproductive success. First, we found that stalk height correlated positively with floral pistil number but not stamen number, suggesting that plants investing in taller stalks benefit more through their seeds than through their pollen. Second, and consistent with this view, the stalks of purely male flowers were shorter than hermaphroditic flowers at all development stages assessed. Third, the positive correlation we observed between stalk height and vegetation height during late but not early season is also consistent with the idea that taller stalks likely improve seed dispersal distance, especially in wind-dispersed species in grassland (Greene and Johnson, 1989; Soons et al., 2004). We found both that larger plants produced taller stalks and that simulated herbivory caused plants to produce shorter stalks in the following season, suggesting that stalk growth represents a net drain on a plant’s resources. In this sense, the cost of floral stalks in *P. alpina* should probably be considered largely as a component of female allocation. Our conjecture that floral stalk growth in *P. alpina* is the outcome of selection via female function more than via male function contrasts with the conclusions reached in studies of other species in which greater floral height more typically enhances pollen export (e.g., O’Connell and Johnston, 1998; Maad, 2000; Eppley and Pannell, 2007), or even that stalk height is under negative selection via male fitness.

The differences in inferred selection via female versus male reproductive success in *P. alpina* may help to explain its ‘andromonoecious’ floral strategy, whereby individuals produce both male and hermaphroditic flowers. Hypotheses for the evolution of andromonoecy have hitherto invoked selection via male reproductive success (Bertin, 1982; Spalik, 1991), with empirical studies asking whether male flowers increase pollinator attraction (Podolsky, 1992; Ehrlén, 1993), enhance pollen output (Elle and Meagher, 2000; Cuevas and Polito, 2004; Dai and Galloway, 2012), or reduce sexual interference within a flower (Quesada-Aguilar et al., 2008). These explanations are inadequate to explain andromonoecy in *P. alpina*. Rather, the association between female sex allocation and stalk height in *P. alpina* constitutes indirect support for an alternative idea, advanced by Jong et al. (2008), that andromonoecy should be favored when the costs of the female function are higher than the costs of stamens and pollinator attraction. This is because the production of both male and hermaphrodite flowers allows plants substantial flexibility to adjust their sex allocation to promote seed dispersal by wind while responding to constraints imposed by their resource status (Spalik, 1991; Zhang and Jiang, 2002): small plants, or those with a resource pool compromised by herbivory or previously high investment in reproduction, reduce their investment in female function (and the necessity to promote it with long stalks) by producing male flowers. Interestingly, the link between wind dispersal of seeds and andromonoecy has been found in other grassland species that also produce elongated stalks, such as species in the Apiaceae (Jury, 1996) and Liliaceae (Peruzzi et al., 2012; Zhang et al., 2014; Niu et al., 2017). It will be important to evaluate these conjectures for *P. alpina* with reference to estimates of both the male and the female components of reproductive success simultaneously.

### Conclusions

Taken together, our results indicate that selection on the ‘simple’ trait of floral stalk height is under the complex influence of both pollinators and seed predators in the context of variation in the sex allocation of the flowers, the height of local competitors in the surrounding vegetation, and the action of defoliating herbivores. The andromonoecious sexual system of *P. alpina* may thus profitably be viewed as a reaction norm of sex allocation at the flower and plant level to these complex interactions. Nevertheless, our study has focused on assessing fitness through only the female component of plant fitness, and a complete picture of selection on reaction norms of sex allocation and the expression of ancillary traits such as floral stalk height awaits a complimentary assessment of selection through male fitness, as well as the mating system, interactions between seed predators and their potential antagonists such as parasitoids, and interactions between genotypes and the environment.

## ACKNOWLEDGEMENTS

This study was funded by a grant from the University of Lausanne. We thank the Canton of Vaud and the Communes of Bex and Ormont-Dessous for access to field sites; Danaé Bataillard and Nora Khelidj for help with fieldwork; and Étienne Lacroix-Carignan and Apiha Shanmuganathan for help with data collection; and members of the Pannell lab for valuable discussions.

## AUTHOR CONTRIBUTION

KS: conceptualization; data curation; formal analysis; methodology; visualization; writing – original draft; writing – review & editing. JRP: funding acquisition; project administration; resources; supervision; validation; writing – review & editing.

## DATA AVAILABILITY

Datasets and codes used in this study were archived in Zenodo (DOI: 10.5281/zenodo.6726264) and can be openly accessed.

## SUPPORTING INFORMATION

Additional supporting information may be found online in the Supporting Information section at the end of the article

Appendix S1. Pollinator assemblage of *P. alpina*.

Appendix S2. Details of studied populations.

Appendix S3. Pictures of *P. alpina* in the field and different categories of achenes.

Appendix S4. Results of the linear mixed model assessing the effects of the intrinsic and extrinsic factors on stalk height at the end of the growing season across eleven populations.

Appendix S5. Results of the linear mixed model assessing the effects of leaf removal treatment and population on stalk height.

Appendix S6. The mean and standard deviation of the four female fitness components and stalk height of the four populations.

Appendix S7. Plots of the single (generalized) linear mixed models assessing the general effects of standardized stalk height on female fitness components.

Appendix S8. Visualization of the relationship between standardized stalk height at the end of the flowering stage and the four fitness components among populations from single (generalized) regression models evaluating the selection gradients without extreme phenotypes.

Appendix S9. Results of the multiple (generalized) linear mixed models assessing the partial effects of four floral traits and population on female fitness components.

## LITERATURE CITED

Ågren, J., F. Hellström, P. Toräng, and J. Ehrlén. 2013. Mutualists and antagonists drive among-population variation in selection and evolution of floral display in a perennial herb. PNAS 110: 18202–18207.

Bates, D., M. Mächler, B. M. Bolker, and S. C. Walker. 2015. Fitting linear mixed-effects models using lme4. Journal of Statistical Software 67: 1–48.

Bertin, R. I. 1982. The evolution and maintenance of andromonoecy. Evolutionary theory 6: 25–32.

Briscoe Runquist, R. D., M. A. Geber, M. Pickett-Leonard, and D. A. Moeller. 2017. Mating system evolution under strong pollen limitation: Evidence of disruptive selection through male and female fitness in Clarkia xantiana. American Naturalist 189: 549–563.

Brodie, E. D., A. J. Moore, and F. J. Janzen. 1995. Visualizing and quantifying natural selection. Trends in Ecology & Evolution 10: 313–318.

Cariveau, D., R. E. Irwin, A. K. Brody, L. S. Garcia-Mayeya, and A. Von Der Ohe. 2004. Direct and indirect effects of pollinators and seed predators to selection on plant and floral traits. Oikos 104: 15–26.

Carlson, J. E., and K. E. Holsinger. 2010. Natural selection on inflorescence color polymorphisms in wild Protea populations: The role of pollinators, seed predators, and intertrait correlations. American Journal of Botany 97: 934–944.

Caruso, C. M., K. E. Eisen, R. A. Martin, and N. Sletvold. 2019. A meta-analysis of the agents of selection on floral traits. Evolution 73: 4–14.

Conner, J. K., S. Rush, S. Kercher, and P. Jennetten. 1996. Measurements of natural selection on floral traits in wild radish (Raphanus raphanistrum). II. Selection through lifetime male and total fitness. Evolution 50: 1137–1146.

Cuevas, J., and V. S. Polito. 2004. The role of staminate flowers in the breeding system of Olea europaea (Oleaceae): An andromonoecious, wind-pollinated taxon. Annals of Botany 93: 547–553.

Dai, C., and L. F. Galloway. 2012. Male flowers are better fathers than hermaphroditic flowers in andromonoecious Passiflora incarnata. New Phytologist 193: 787–796.

Diniz, U. M., A. Domingos-Melo, and I. C. MacHado. 2019. Flowers up! The effect of floral height along the shoot axis on the fitness of bat-pollinated species. Annals of Botany 124: 809–818.

Donohue, K. 1998. Maternal determinants of seed dispersal in Cakile edentula: Fruit, plant, and site traits. Ecology 79: 2771–2788.

Donohue, K. 1999. Seed dispersal as a maternally influenced character: Mechanistic basis of maternal effects and selection on maternal characters in an annual plant. American Naturalist 154: 674–689.

Ehrlén, J. 1993. Ultimate functions of non-fruiting flowers in Lathyrus vernus. Oikos 68: 45.

Ehrlén, J., A.-K. Borg-Karlson, and A. Kolb. 2012. Selection on plant optical traits and floral scent: Effects via seed development and antagonistic interactions. Basic and Applied Ecology 13: 509–515.

Elle, E., and T. R. Meagher. 2000. Sex allocation and reproductive success in the andromonoecious perennial Solanum carolinense (Solanaceae). II. Paternity and functional gender. American Naturalist 156: 622–636.

Eppley, S. M., and J. R. Pannell. 2007. Density-dependent self-fertilization and male versus hermaphrodite siring success in an androdioecious plant. Evolution 61: 2349–2359.

Frey, F. M. 2007. Opposing natural selection from herbivores and pathogens may maintain floral-color variation in Claytonia virginica (Portulacaceae). Evolution 58: 2426–2437.

Galen, C. 1989. Measuring pollinator-mediated selection on morphometric floral traits: bumblebees and the alpine sky pilot, Polemonium viscosum. Evolution 43: 882–890.

Giunti, G., A. Canale, R. H. Messing, E. Donati, C. Stefanini, J. P. Michaud, and G. Benelli. 2015. Parasitoid learning: Current knowledge and implications for biological control. Biological Control 90: 208–219.

Gómez, J. M. 2007. Dispersal-mediated selection on plant height in an autochorously dispersed herb. Plant Systematics and Evolution 268: 119–130.

Gómez, J. M. 2003. Herbivory reduces the strength of pollinator-mediated selection in the mediterranean herb Erysimum mediohispanicum: Consequences for plant specialization. American Naturalist 162: 242–256.

Gómez, J. M. 2008. Sequential conflicting selection due to multispecific interactions triggers evolutionary trade-offs in a monocarpic herb. Evolution; international journal of organic evolution 62: 668–679.

Gómez, J. M., F. Perfectti, J. Bosch, and J. P. M. Camacho. 2009. A geographic selection mosaic in a generalized plant-pollinator-herbivore system. Ecological Monographs 79: 245–263.

Gómez, J. M., and R. Zamora. 2000. Spatial variation in the selective scenarios of Hormathophylla spinosa (Cruciferae). American Naturalist 155: 657–668.

Gómez, J. M., and R. Zamora. 1994. Top-down effects in a tritrophic system: Parasitoids enhance plant fitness. Ecology 75: 1023–1030.

Greene, D. F., and E. A. Johnson. 1989. A model of wind dispersal of winged or plumed seeds. Ecology 70: 339–347.

Harder, L. D., and P. Prusinkiewicz. 2013. The interplay between inflorescence development and function as the crucible of architectural diversity. Annals of Botany 112: 1477–1493.

Hartig, F. 2019. DHARMa: residual diagnostics for hierarchical (multi-level/mixed) regression models. R package version 0.2.4.

Hodgins, K. A., and S. C. H. Barrett. 2008. Natural selection on floral traits through male and female function in wild populations of the heterostylous daffodil Narcissus triandrus. Evolution 62: 1751–1763.

Janzen, F. J., and H. S. Stern. 1998. Logistic regression for empirical studies of multivariate selection. Evolution 52: 1564–1571.

Jervis, M. A., and N. A. C. Kidd. 1986. Host-feeding strategies in Hymenopteran parasitoids. Biological Reviews 61: 395–434.

de Jong, T. J., and P. G. L. Klinkhamer. 1989. Size-dependency of sex-allocation in hermaphroditic, monocarpic plants. Functional Ecology 3: 201–206.

de Jong, T. J., A. Shmida, and F. Thuijsman. 2008. Sex allocation in plants and the evolution of monoecy. Evolutionary Ecology Research 10: 1087–1109.

Jury, S. L. 1996. Pollination and dispersal in Mediterranean umbellifers. Bocconea 5: 193–199.

Kessler, D., M. Kallenbach, C. Diezel, E. Rothe, M. Murdock, and I. T. Baldwin. 2015. How scent and nectar influence floral antagonists and mutualists. eLife 4: e07641.

Klecka, J., J. Hadrava, P. Biella, and A. Akter. 2018. Flower visitation by hoverflies (Diptera: Syrphidae) in a temperate plant-pollinator network. PeerJ 6: e6025.

Klinkhamer, P., T. de Jong, and H. Metz. 1997. Sex and size in cosexual plants. Trends in Ecology & Evolution 12: 260–265.

Knauer, A. C., and F. P. Schiestl. 2017. The effect of pollinators and herbivores on selection for floral signals: a case study in Brassica rapa. Evolutionary Ecology 31: 285–304.

Lande, R., and S. J. Arnold. 1983. The measurement of selection on correlated characters. Evolution 37: 1210–1226.

Lauber, K., G. Wagner, A. Gygax, S. Eggenberg, and A. Michel. 2018. Flora helvetica. 5th ed. Haupt Bern.

Lenth, R. V. 2020. Emmeans: Estimated marginal means, aka least-squares means.

Maad, J. 2000. Phenotypic selection in hawkmoth-pollinated Plntathera bifolia: Targets and fitness surfaces. Evolution 54: 112–123.

Matsumura, S., R. Arlinghaus, and U. Dieckmann. 2012. Standardizing selection strengths to study selection in the wild: A critical comparison and suggestions for the future. BioScience 62: 1039–1054.

Molau, U., B. Eriksen, and J. T. Knudsen. 1989. Predispersal seed predation in Bartsia alpina. Oecologia 81: 181–185.

Morrissey, M. B. 2014. In search of the best methods for multivariate selection analysis. Methods in Ecology and Evolution 5: 1095–1109.

Muller-Schneider, P. 1986. Verbreitungsbiologie der Blutenpflanzen Graubundens. Veroffentlichungen des geobotanischen Institutes der eidgenossischen technischen Hochschule, Stiftung Rubel 85: 261.

Niu, Y., Q. Gong, D. Peng, H. Sun, and Z. Li. 2017. Function of male and hermaphroditic flowers and size-dependent gender diphasy of Lloydia oxycarpa (Liliaceae) from Hengduan Mountains. Plant Diversity 39: 187–193.

O’Connell, L. M., and M. O. Johnston. 1998. Male and female pollination success in a deceptive orchid, a selection study. Ecology 79: 1246–1260.

Parachnowitsch, A. L., J. S. Manson, and N. Sletvold. 2019. Evolutionary ecology of nectar. Annals of Botany 123: 247–261.

Pérez-Barrales, R., G. H. Bolstad, C. Pélabon, T. F. Hansen, and W. S. Armbruster. 2013. Pollinators and seed predators generate conflicting selection on Dalechampia blossoms. Oikos 122: 1411–1428.

Peruzzi, L., E. Mancuso, and D. Gargano. 2012. Males are cheaper, or the extreme consequence of size/age-dependent sex allocation: sexist gender diphasy in Fritillaria montana (Liliaceae). Botanical Journal of the Linnean Society 168: 323–333.

Pickup, M., and S. C. H. Barrett. 2012. Reversal of height dimorphism promotes pollen and seed dispersal in a wind-pollinated dioecious plant. Biology Letters 8: 245–248.

Podolsky, R. D. 1992. Strange floral attractors: Pollinator attraction and the evolution of plant sexual systems. Science 258: 791–793.

Quesada-Aguilar, A., S. Kalisz, and T. L. Ashman. 2008. Flower morphology and pollinator dynamics in Solanum carolinense (Solanaceae): Implications for the evolution of andromonoecy. American Journal of Botany 95: 974–984.

R Core Team. 2021. R: a language and environment for statistical computing.

Schiestl, F. P., F. K. Huber, and J. M. Gomez. 2011. Phenotypic selection on floral scent: Trade-off between attraction and deterrence? Evolutionary Ecology 25: 237–248.

Schiestl, F. P., and S. D. Johnson. 2013. Pollinator-mediated evolution of floral signals. Trends in Ecology & Evolution 28: 307–315.

Sletvold, N., and J. Ågren. 2015. Pollinator-mediated selection on floral display and spur length in the orchid Gymnadenia conopsea. International Journal of Plant Sciences 171: 999–1009.

Sletvold, N., J. M. Grindeland, and J. Agren. 2013. Vegetation context influences the strength and targets of pollinator-mediated selection in a deceptive orchid. Ecology 94: 1236–1242.

Sletvold, N., K. K. Moritz, and J. Ågren. 2015. Additive effects of pollinators and herbivores result in both conflicting and reinforcing selection on floral traits. Ecology 96: 214–221.

Soons, M. B., G. W. Heil, R. Nathan, and G. G. Katul. 2004. Determinants of long-distance seed dispersal by wind in grasslands. Ecology 85: 3056–3068.

Spalik, K. 1991. On evolution of andromonoecy and ‘overproduction’ of flowers: A resource allocation model. Biological Journal of the Linnean Society 42: 325–336.

Stinchcombe, J. R., A. F. Agrawal, P. A. Hohenlohe, S. J. Arnold, and M. W. Blows. 2008. Estimating nonlinear selection gradients using quadratic regression coefficients: Double or nothing? Evolution 62: 2435–2440.

Strauss, S. Y., and J. B. Whittall. 2006. Non-pollinator agents of selection on floral traits. In L. D. Harder, and S. C. H. Barrett [eds.], Ecology and Evolution of Flowers, 120–138. Oxford University Press.

Szentpéteri, J. L., S. Szente, B. Szego, and S. Stranczinger. 2008. Morphological studies and taxonomic review of Preonanthus (Ranunculaceae). Acta Botanica Hungarica 50: 407–415.

Thomann, M., J. Ehrlén, and J. Ågren. 2018. Grazers affect selection on inflorescence height both directly and indirectly and effects change over time. Ecology 99: 2167–2175.

Thompson, J. N. 1999. Specific hypotheses on the geographic mosaic of coevolution. American Naturalist 153: S1–S14.

Tonnabel, J., P. David, E. K. Klein, and J. R. Pannell. 2019. Sex-specific selection on plant architecture through “budget” and “direct” effects in experimental populations of the wind-pollinated herb, Mercurialis annua. Evolution 73: 897–912.

Vittoz, P., and R. Engler. 2007. Seed dispersal distances: A typology based on dispersal modes and plant traits. Botanica Helvetica 117: 109–124.

Waite, S., and M. J. Hutchings. 1982. Plastic energy allocation patterns in Plantago coronopus. Oikos 38: 333.

Waser, N. 1983. The adaptive nature of floral traits: Ideas and evidence. In L. Real [ed.], Pollination Biology, 241–285. Academic Press.

Weiner, J. 2004. Allocation, plasticity and allometry in plants. Perspectives in Plant Ecology, Evolution and Systematics 6: 207–215.

Winkler, I. S., S. J. Scheffer, and C. Mitter. 2009. Molecular phylogeny and systematics of leaf-mining flies (Diptera: Agromyzidae): Delimitation of Phytomyza Fallén sensu lato and included species groups, with new insights on morphological and host-use evolution. Systematic Entomology 34: 260–292.

Zhang, D. Y., and X. H. Jiang. 2002. Size-dependent resource allocation and sex allocation in herbaceous perennial plants. Journal of Evolutionary Biology 15: 74–83.

Zhang, Z. Q., X. F. Zhu, H. Sun, Y. P. Yang, and S. C. H. Barrett. 2014. Size-dependent gender modification in Lilium apertum (Liliaceae): Does this species exhibit gender diphasy? Annals of Botany 114: 441–453.

