## Appendix S1 for "Disruptive selection via pollinators and seed predators on the height of flowers in a wind-dispersed alpine herb"

**Appendix S1.** Pollinator assemblage of *P. alpina* in population LM in 2018. The frequency of different groups of insect visitors during the course of the flowering season. Three to five 10-minute observations were conducted on each observation day. All the insects observed visiting flowers were recorded. In total, 265 floral visitations were recorded. Hymenoptera, Lepidoptera, and Hemiptera visitors are shown in the category “Others”. Diptera, including the house fly and syrphid flies, were the dominant floral visitors of *P. alpina*.

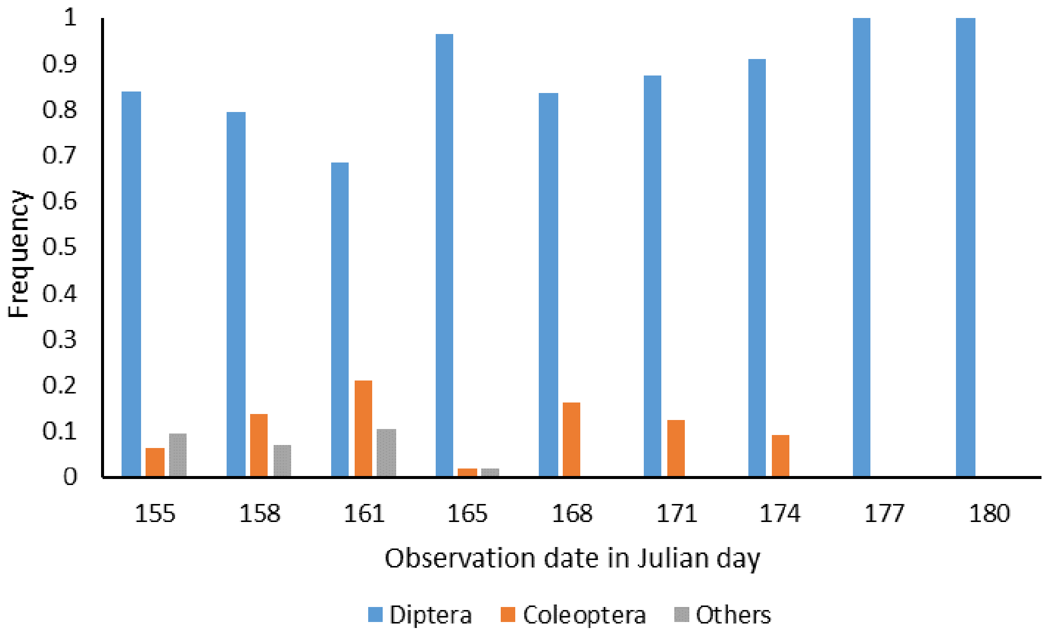
