## Appendix S2 for "Disruptive selection via pollinators and seed predators on the height of flowers in a wind-dispersed alpine herb"

### **Appendix S2.** Details of studied populations

* Flowering season indicates whether the population starts flowering in late May to early June (E) or late June to July (L).

+ Grazing is quantified in terms of three categories, based on personal observation in 2019, i.e., no grazing (0), highly grazed (1), and intermediate level (0.5).

[!] Measurement was not conducted in LL1 population due to a landslide in 2020 and in S1 population due to unintended herbivory at the end of the growing season in 2021.

| Population | Location | GPS position | Altitude (a.s.l.) | Flower season * | Grazing [+] | Habitat | Note | Population survey[!] (2021) | Floral correlation (2021) | Leaf removal (2020-2021) | Selection gradient (2019) |
|---|---|---|---|---|---|---|---|---|---|---|---|
| S1 | Solalex | 46°17′36N 7°09′11″E | 1723 | E | 0 | open grassland | fenced | | O | O | O |
| S1+ | Solalex | 46°17′42″N 7°09′09″E | 1758 | E | 1 | open grassland | shallow soil | O | | | |
| S2 | Solalex | 46°16′37″N 7°09′32″E | 2122 | L | 0.5 | open grassland | steep | O | | | O |
| S4 | Solalex | 46°16′42″N 7°09′47″E | 2003 | L | 1 | open grassland | flat, shallow soil | O | | | |
| B1 | Bretaye | 46°19′31″N 7°05′00″E | 1818 | E | 0 | open grassland | | O | | | |
| LS1 | Leysin | 46°21′42″N 7°00′13″E | 1995 | L | 0.5 | open grassland | | O | | | |
| LS2 | Leysin | 46°21′40″N 7°00′15″E | 2006 | E | 0 | open grassland | shallow soil | O | | | |
| R1 | Rochers de Naye | 46°25′58″N 6°59′04″E | 1953 | E | 0 | open grassland | steep, fenced | O | | | |
| R2 | Rochers de Naye | 46°25′58″N 6°59′03″E | 1944 | L | 0.5 | open grassland | | O | | | |
| LM1 | Les Mosses | 46°23′57″N 7°04′52″E | 1694 | E | 1 | enclosed grassland | | O | | | |
| LL1 | Lac Lioson | 46°23′08″N 7°07′23″E | 1951 | E | 0.5 | open grassland | steep | | | | O |
| LL3 | Lac Lioson | 46°23′05″N 7°07′25″E | 1900 | L | 0.5 | open grassland | | O | | O | |
| LL4 | Lac Lioson | 46°22′57″N 7°07′11″E | 1983 | L | 0.5 | open grassland | | O | | | O |
