## Appendix S3 for "Disruptive selection via pollinators and seed predators on the height of flowers in a wind-dispersed alpine herb"

**Appendix S3.** Pictures of *Pulsatilla alpina* in the field and different categories of achenes. **(A)** A flowering individual. **(B)** Elongated stalks bearing achenes before dispersal at the end of the growing season. **(C)** Different categories of achenes. From the top to the bottom is an unfertilized achene, a predated achene, and a mature achene. Top achene: the color of unfertilized achenes is usually pale brown to yellow. The head part is flat and soft. Middle achene: Predated achenes have the same color and size as mature ones but have a hole made by seed predators (indicated by a red triangle). When squeezed, they are soft and empty. Bottom achene: Mature achenes are robust and contain a seed within a brown-dark head capsule. The heads of the achenes are swollen and hard. The white bar in **(C)** indicates one centimeter.

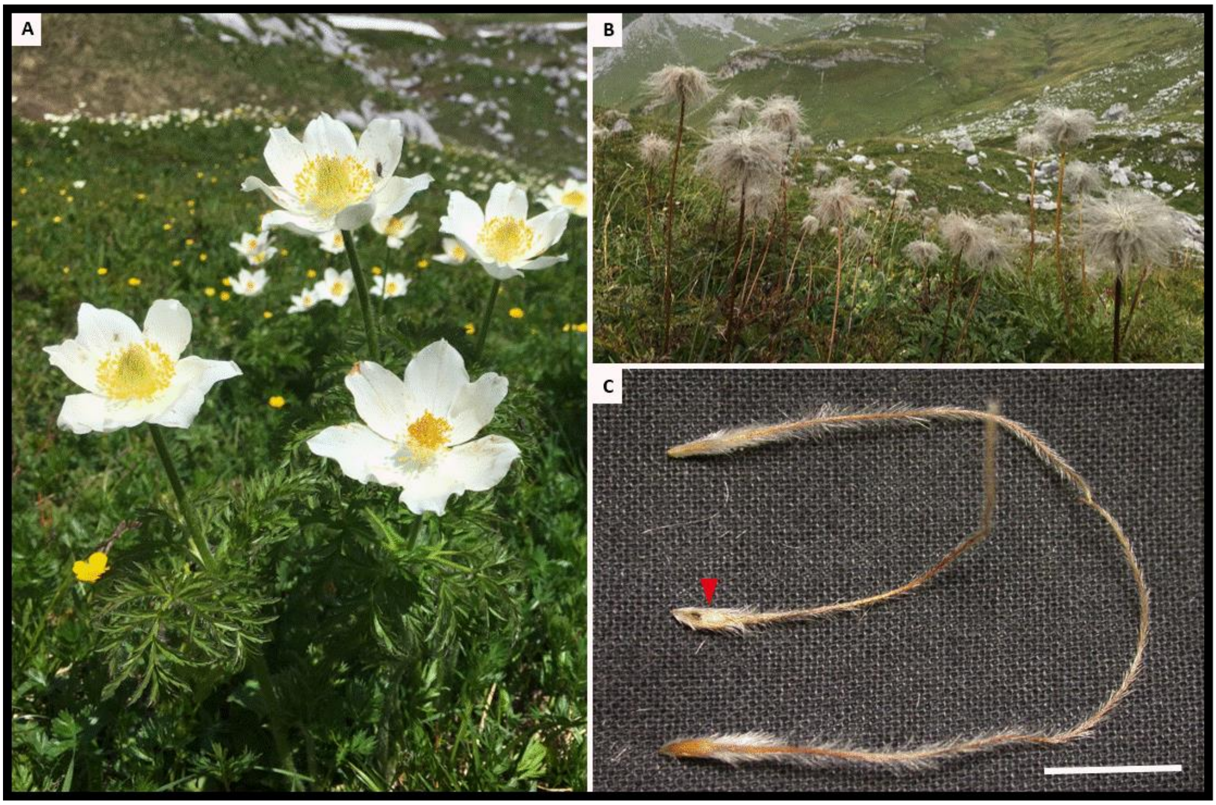
