## Appendix S4 for "Disruptive selection via pollinators and seed predators on the height of flowers in a wind-dispersed alpine herb"

**Appendix S4.** Results of the linear mixed model assessing the effects of the intrinsic and extrinsic factors on stalk height at the end of the growing season across eleven populations.

Notes: ∗ P < 0.05, ∗∗ P < 0.01, ∗∗∗ P < 0.001

| **Fixed effect** | **df** | **Likelihood-ratio test** | ***P*** |
|---|---|---|---|
| Total flower number | 1 | 68.4 | *** |
| Vegetation height at early season | 1 | 0.27 | ns |
| Vegetation height at late season | 1 | 9.66 | ** |
| Population | 10 | 55.0 | *** |
| **Random effect** | **N** | **Variance component** | **SD** |
| Residuals | | 43.7 | 6.61 |
| Transect | 41 | 51.3 | 2.44 |
