## Appendix S5 for "Disruptive selection via pollinators and seed predators on the height of flowers in a wind-dispersed alpine herb"

**Appendix S5.** Results of the linear mixed model assessing the effects of leaf removal treatment and population on stalk height.

Notes: ∗ $P < 0.05$, ∗∗ $P < 0.01$, ∗∗∗ $P < 0.001$

| **Fixed effect** | **df** | **Likelihood-ratio test** | ***P*** |
|---|---|---|---|
| Leaf removal | 1 | 9.05 | ** |
| Population | 1 | 5.02 | * |
| Leaf removal: Population | 1 | 0.83 | ns |
| **Random effect** | **N** | **Variance component** | **SD** |
| Residuals | | 51.3 | 7.16 |
| Individual | 63 | 50.7 | 7.12 |
