## Appendix S6 for "Disruptive selection via pollinators and seed predators on the height of flowers in a wind-dispersed alpine herb"

**Appendix S6.** The mean and standard deviation of the four female fitness components and stalk height of the four populations. The mean and standard deviation were calculated at the flower-level in the four populations: S1 (N = 95 flowers), S2 (N = 74 flowers), LL1 (N = 62 flowers), and LL4 (N = 91 flowers).

| Population | Fertilization rate | Rate of non-predation | Seed maturation rate | Mature seed number | Stalk height |
|---|---|---|---|---|---|
| LL1 | 0.37 ± 0.18 | 0.71 ± 0.22 | 0.27 ± 0.16 | 51.2 ± 35.9 | 43.7 ± 6.68 |
| LL4 | 0.60 ± 0.16 | 0.53 ± 0.20 | 0.33 ± 0.17 | 72.2 ± 42.9 | 45.9 ± 7.69 |
| S1 | 0.53 ± 0.21 | 0.66 ± 0.21 | 0.38 ± 0.20 | 71.8 ± 45.9 | 43.5 ± 7.51 |
| S2 | 0.49 ± 0.19 | 0.59 ± 0.22 | 0.30 ± 0.17 | 60.0 ± 36.9 | 36.9 ± 7.18 |
