## Appendix S7 for "Disruptive selection via pollinators and seed predators on the height of flowers in a wind-dispersed alpine herb"

**Appendix S7.** Plots of the single (generalized) linear mixed models assessing the general effects of standardized stalk height on female fitness components. The relationship between stalk height at the end of the flowering stage and **(A)** fertilization rate, **(B)** rate of non-predation, **(C)** seed maturation rate, and **(D)** relative mature achene number from single regression models. Grey points are flowers from the four populations studied in 2019. Interactions between stalk height and population are all non-significant. The shaded ribbon indicates the 95% confidence interval of the regression lines.

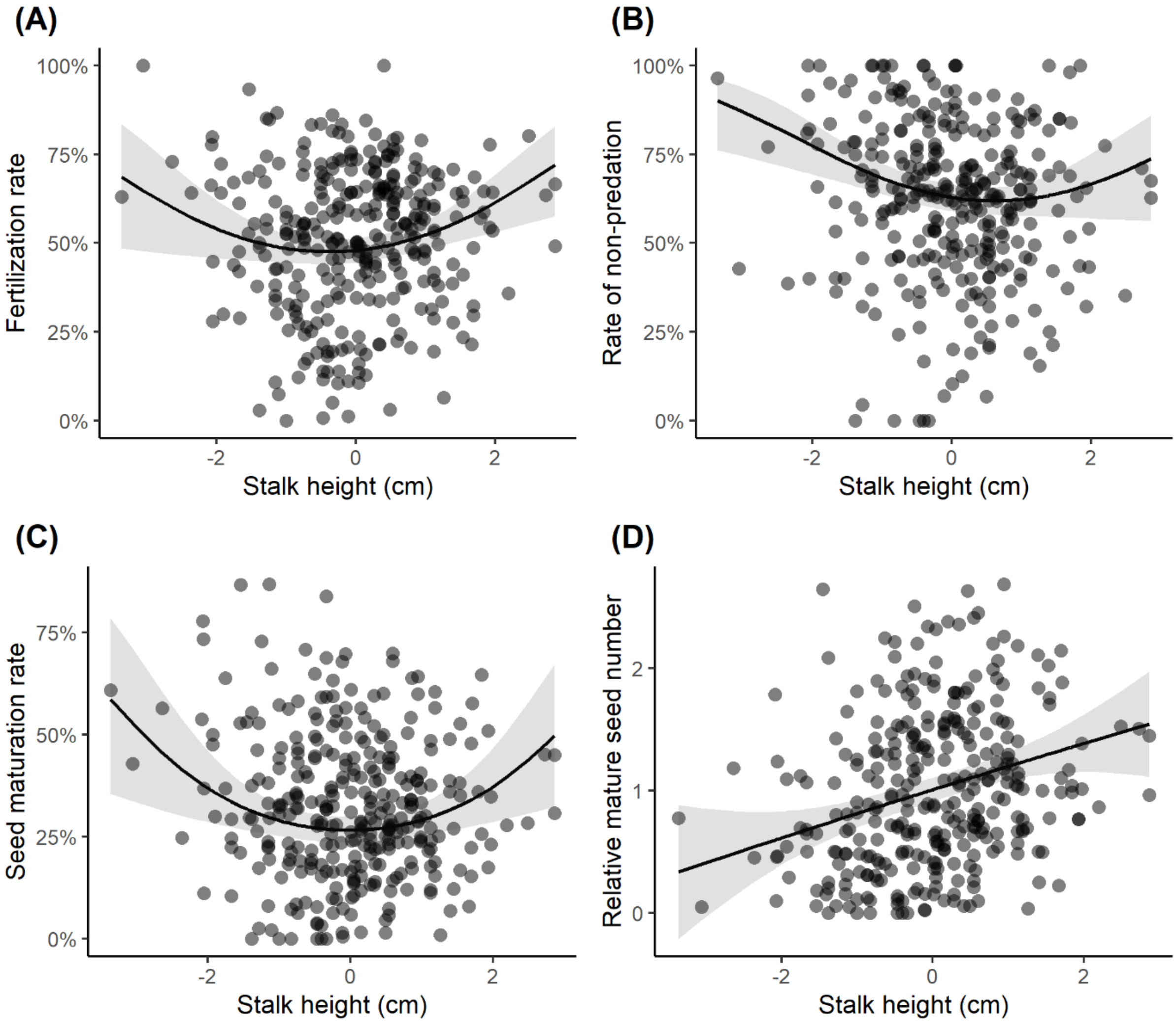
