## Appendix S8 for "Disruptive selection via pollinators and seed predators on the height of flowers in a wind-dispersed alpine herb"

**Appendix S8.** Visualization of the relationship between standardized stalk height at the end of the flowering stage and (A) fertilization rate, (B) rate of non-predation, (C) seed maturation rate, and (D) relative mature seed number among populations from single regression models evaluating the selection differentials without extreme phenotypes (i.e., phenotypes larger or small than two standard deviations from the mean value of each population; 14 data points removed from a total of 322 data points). Raw data points and regression lines of the studied populations are shown in different colors. The shaded ribbons indicate the standard error of the regression lines. The results indicate that the selection differentials and gradients estimated might be largely driven by those extreme phenotypes. However, it seems the extreme phenotypes have shaped the selection in the same direction in all four populations (see Fig. 4 for a comparison). The linear and quadratic differentials for fertilization rate, rate of non-predation, and seed maturation rate were non-significant in all the populations except the quadratic coefficients in LL1 population ($P < 0.001$, $P = 0.05$, and $P < 0.01$ in the three fitness components, respectively). For relative mature seed number, the quadratic coefficients were all non-significant, while the linear coefficients were all significant except for LL1 (marginally n.s., $P = 0.06$) and LL4 (n.s., $P = 0.18$) population. Stronger approaches one can use in further studies to overcome the issue are either to manipulate and create extreme phenotypes or to select extreme phenotypes from the natural variation to have a better estimate of the selection at the two ends of the phenotypic range.

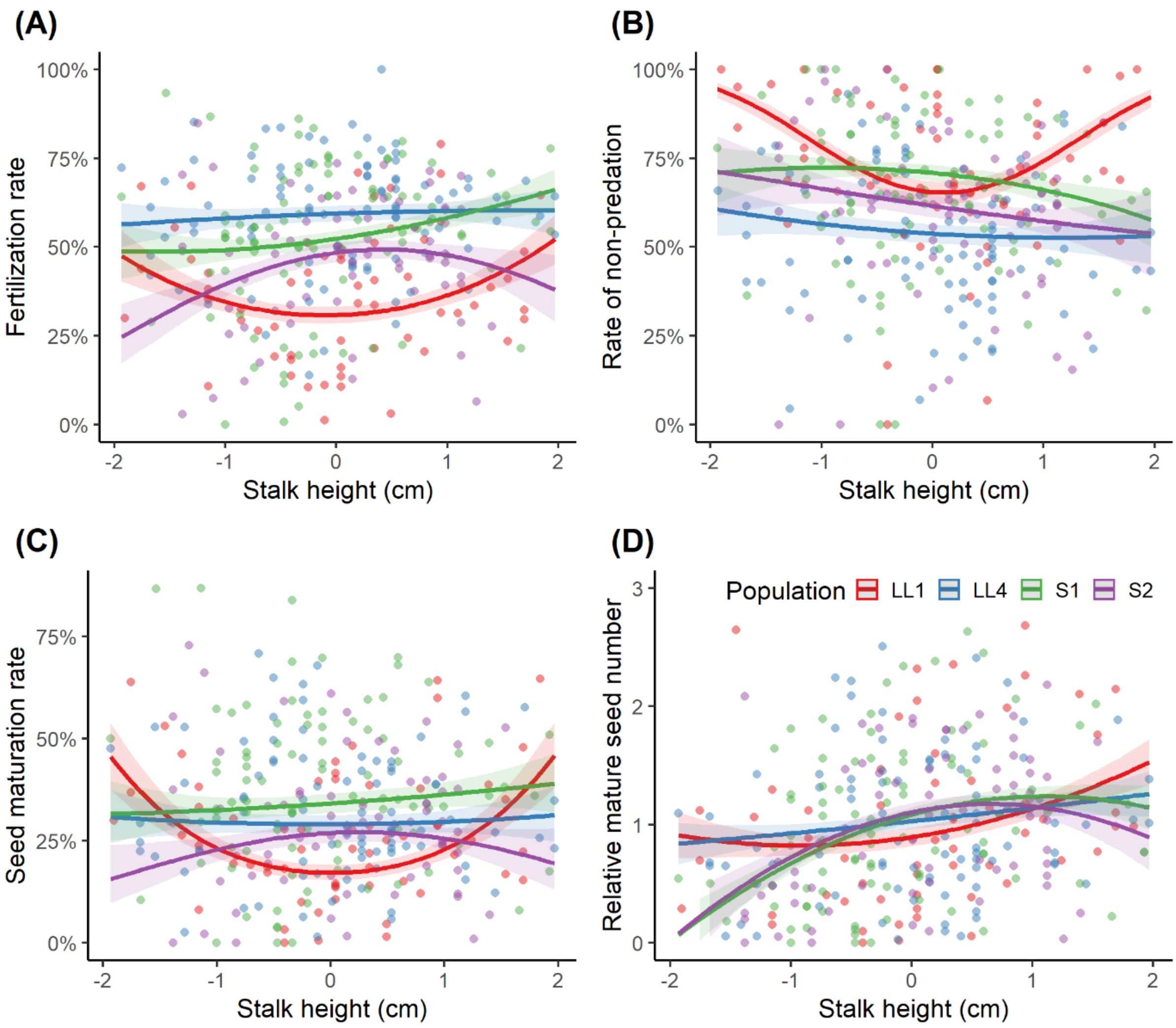

(A)
Fertilization rate
100%
75%
50%
25%
0%
-2
-1
0
1
2
Stalk height (cm)
(B)
Rate of non-predation
100%
75%
50%
25%
0%
-2
-1
0
1
2
Stalk height (cm)
(C)
Seed maturation rate
75%
50%
25%
0%
-2
-1
0
1
2
Stalk height (cm)
(D)
Population
LL1
LL4
S1
S2
Relative mature seed number
3
2
1
0
-2
-1
0
1
2
Stalk height (cm)
