## Appendix S9 for "Disruptive selection via pollinators and seed predators on the height of flowers in a wind-dispersed alpine herb"

**Appendix S9.** Results of the generalized linear mixed models assessing the effects of four floral traits and population on female fitness components.

Notes: ∗ P < 0.05, ∗∗ P < 0.01, ∗∗∗ P < 0.001

| | Response variable | | | | | | | | | | | |
|---|---|---|---|---|---|---|---|---|---|---|---|---|
| | **Fertilization rate (binomial)** | | | **Rate of non-predation (binomial)** | | | **Seed maturation rate (binomial)** | | | **Relative mature seed number (normal)** | | |
| **Fixed effect** | **df** | **LRT** | ***P*** | **df** | **LRT** | ***P*** | **df** | **LRT** | ***P*** | **df** | **LTR** | ***P*** |
| Stalk height | 1 | 0.44 | ns | 1 | 0.00 | ns | 1 | 0.19 | ns | 1 | 0.26 | ns |
| Tepal length | 1 | 5.58 | * | 1 | 1.48 | ns | 1 | 7.26 | ** | 1 | 4.32 | * |
| Pistil number | 1 | 0.00 | ns | 1 | 30.0 | *** | 1 | 6.84 | ** | 1 | 85.6 | *** |
| Stamen number | 1 | 9.88 | ** | 1 | 3.69 | . | 1 | 12.5 | *** | 1 | 5.04 | * |
| Stalk height$^2$ | 1 | 5.86 | * | 1 | 3.90 | * | 1 | 7.79 | ** | 1 | 2.57 | ns |
| Tepal length$^2$ | 1 | 0.57 | ns | 1 | 0.07 | ns | 1 | 0.37 | ns | 1 | 0.46 | ns |
| Pistil number$^2$ | 1 | 0.43 | ns | 1 | 1.86 | ns | 1 | 1.55 | ns | 1 | 9.43 | ** |
| Stamen number$^2$ | 1 | 4.53 | * | 1 | 0.05 | ns | 1 | 2.25 | ns | 1 | 2.26 | ns |
| Population | 3 | 35.1 | *** | 3 | 21.5 | *** | 3 | 9.40 | * | 3 | 0.24 | ns |
| Stalk height: Population | 3 | 0.54 | ns | 3 | 6.73 | . | 3 | 1.13 | ns | 3 | 2.04 | ns |
| Tepal length: Population | 3 | 1.47 | ns | 3 | 4.63 | ns | 3 | 2.18 | ns | 3 | 1.61 | ns |
| Pistil number: Population | 3 | 1.58 | ns | 3 | 13.6 | ** | 3 | 3.08 | ns | 3 | 1.37 | ns |
| Stamen number: Population | 3 | 1.83 | ns | 3 | 4.90 | ns | 3 | 4.81 | ns | 3 | 5.30 | ns |
| Stalk height$^2$: Population | 3 | 3.14 | ns | 3 | 6.71 | . | 3 | 2.44 | ns | 3 | 3.24 | ns |
| Tepal length$^2$: Population | 3 | 3.17 | ns | 3 | 3.70 | ns | 3 | 4.28 | ns | 3 | 3.59 | ns |
| Pistil number$^2$: Population | 3 | 0.92 | ns | 3 | 0.84 | ns | 3 | 1.90 | ns | 3 | 9.86 | ** |
| Stamen number$^2$: Population | 3 | 4.59 | ns | 3 | 3.80 | ns | 3 | 6.54 | . | 3 | 3.73 | ns |
| **Random effect** | **N** | **Variance component** | **SD** | **N** | **Variance component** | **SD** | **N** | **Variance component** | **SD** | **N** | **Variance component** | **SD** |
| Flower ID | 322 | 0.52 | 0.72 | 321 | 0.65 | 0.81 | 322 | 0.63 | 0.79 | | | |
| Individual ID | 151 | 0.14 | 0.37 | 151 | 0.21 | 0.46 | 151 | 0.22 | 0.47 | 151 | 0.09 | 0.29 |
